# Precision Prediction of Microbial Ecosystem Impact on Host Metabolism Using Genome-Resolved Metagenomics

**DOI:** 10.1101/2025.06.27.661827

**Authors:** Mohamed Mohssen, Ahmed A. Zayed, Kristina A. Kigerl, Jingjie Du, Garrett J. Smith, Jan M. Schwab, Matthew B. Sullivan, Phillip G. Popovich

**Affiliations:** The Interdisciplinary Biophysics Graduate Program, The Ohio State University, Columbus, Ohio 43210, USA; Department of Microbiology, The Ohio State University, Columbus, Ohio 43210, USA; Center of Microbiome Science, The Ohio State University, Columbus, Ohio 43210, USA; EMERGE Biology Integration Institute, The Ohio State University, Columbus, Ohio 43210, USA; Department of Neuroscience, The Ohio State University Wexner Medical Center, Columbus, OH, 43210, USA; Belford Center for Spinal Cord Injury, The Ohio State University Wexner Medical Center, Columbus, OH, 43210, USA; Department of Civil, Environmental and Geodetic Engineering, The Ohio State University, Columbus, OH 43210, USA

## Abstract

Microorganisms often drive ecosystem function, yet precision disturbance response and ecosystem impact predictions remain challenging due to poorly captured ecological and metabolic interconnectedness and functional redundancy. For example, while mammalian gut dysbiosis is recognized to influence host metabolism, key microbiota and mechanisms governing their effects remain poorly understood. Here we developed a genome-resolved eco-systems biology workflow to predict how gut microbial metabolism affects mammalian health, and we applied it to a ‘spinal cord-gut axis’ dataset. By scaling and integrating temporally resolved network analytics and consensus statistical approaches, we identified largely previously uncharacterized microbial species that best predict host physiology following neurological impairment. *In silico* validation through “complete” pathway-centric and comparative genomic analyses revealed that among these species, the major encoded microbial metabolic changes were in pathways directly linked to host nitrogen balance, and they varied by host sex and microbial ecotype/species. Moreover, we identified the exact bacterial species (and their draft genome sequences) driving urease-dependent versus amino acid-dependent nitrogen gut metabolism – findings that explain previously mechanistically-ambiguous, but clinically relevant, ammonia-driven host nitrogen imbalance. More broadly, these ecology- and community-aware approaches provide a framework to study dynamic, interconnected microbiomes that advances from enrichment-based single-taxon and single-gene correlations towards building microbe(s)-driven mechanistic insights that integrate community context and whole pathways.

## Introduction

Earth systems are inhabited by microorganisms that perform key elemental transformations, including those related to carbon^1,2^ and nitrogen^3^ cycling. In these systems, ecological context is critical for understanding microbiota functioning, especially when it comes to microbial adaptations to particular niches^4^. While gene marker approaches provide incredibly valuable microbial community composition data for new ecosystems studied, genome-resolved approaches are emerging as the gold standard where it is critical to capture the fast-evolving, niche-defining functions that dictate ecosystem functioning and that marker genes miss^5–14^, particularly essential for translating the microbiome in health and disease^13,15–17^. Additionally, microbiomes are dynamic and highly interconnected as microbes exchange and compete for metabolites and other molecules in their environment^18,19^ (e.g., the gut^20^).

Despite the growing appreciation of microbial contributions to gut ecosystem function, current approaches to study these systems largely focus on microbiota constituents in isolation (i.e., linking a single taxon/gene at a time to health/disease), often ignoring complex and dynamic whole community interactions. Recently, specific microbes have been identified to significantly impact health and disease, including *Faecalibacterium* species in inflammatory bowel diseases and colorectal cancer (reviewed in^21^), *Hominenteromicrobium* strain YB328 in enhancing anti-tumour immunity^22^, and several others (reviewed in^6,23^). However, such microbes undoubtedly rely on ecosystem-dependent metabolic interactions within the microbial community, with a recently elucidated example being carbon-based metabolic interplay for beneficial *Lactobacillus johnsonii*^24^. Beyond genomic representation at the community level, gut bacterial gene expression profiles are also emerging as taxon-context-dependent^25^. Thus, even if one taxon leads to significant health benefits under an experimental condition(s), ignoring microbe-microbe interactions limits the generalizability and modulation of microbe-mediated interventions. This calls for an eco-systems biology approach^26^ whereby data generation captures genome-wide variation (including ecosystem-specific, niche-defining traits) and scalable, hypothesis-free analytics account for complex functional redundancies and microbial interactions community-wide to best estimate ecosystem functioning.

Here we sought to establish such a “data-driven” workflow to (i) better identify emergent ecosystem properties that elucidate precise microbial contributions to host health and disease and (ii) apply this workflow to a biomedically-relevant dataset as a case study. To do this, we focused on neurological impairment (NI) as a gut ecosystem “environmental” challenge^27–34^ that alters mammalian host metabolism^35,36^. This systematic workflow developed to investigate microbe-host interactions requires no *a priori* assumptions and uses (i) cross-validated eco-systems biology analytics, including machine learning, to identify key ecosystem-impacting microbial taxa, and (ii) consensus genome-resolved, pathway-centric (i.e., requiring multiple genes involved in a pathway/function) analyses to elucidate metabolism-based mechanisms underpinning these ecosystem impacts. These requirements to evaluate enriched functions in community and pathway context take a critical step from pairwise correlations towards systems-level functional assessments of causation and mechanism. Case study application of this data-driven approach highlights key microbial species and metabolic “complete” pathways influencing mammalian host nitrogen metabolism and provides insights into potential therapeutic targets for future *in vivo* validation.

### Emergent gut microbial networks predict host physiology

We previously established a well-controlled spinal cord-gut axis murine model system^24^ that represents NI and includes high-resolution gut metagenome temporal sampling, and we manually reconstructed carbon-based microbial metabolic connectivity in the gut ecosystem to inform microbe-microbe interactions. To evaluate microbe-host metabolic interactions in a whole-ecosystem context, we leverage the curated Mouse B6 Gut Catalog (MB6GC)^24^, which includes 263 dereplicated metagenome-assembled genomes (MAGs) recovered from fecal and cecal samples temporally obtained from male and female mice with graded levels of NI caused by spinal lesions at mid- (T10) or high- (T4) thoracic spinal levels. Such lesions partially or completely disrupt neural control over the gut. Mice receiving spinal surgery, but without a spinal cord lesion, served as a control group (“Lam”) (see^24^ for full study design). These MAGs, which represented the *in-situ* gut microbial community quite well (i.e., they recruited ∼80% of the total metagenomic reads), were quantified over 6 months across groups and evaluated against a standardized scale of neurological function in mice (Basso Mouse Scale, or BMS, for locomotion) as a proxy for overall mouse physiological state.

To analyze these data considering the full microbial community context, we applied an eco-systems biology approach (*sensu*^37^) that uses MAG-based relative abundance data to organize the microbial community into predictably interacting (e.g., competing or cross-feeding) sub-communities of co-occurring MAGs via weighted gene/genome correlation network analysis (WGCNA)^38^. Longitudinal sample-dependencies were accounted for by generating six independent networks – one for each timepoint across 6 months following NI (see **Fig. 1A**, **Fig. S1**, and “*WGCNA Analysis”* in the **Methods** for more details). The resulting initial networks were power-transformed to identify robust subnetworks (i.e., modules) meant to represent microbial sub-communities whose members co-vary and therefore possibly interact. Each subnetwork was then correlated with host neurological function (BMS scores) such that member taxa of correlating subnetworks could be analyzed via leave-one-out cross-validation to identify members critical for these associations. Specifically, leave-one-out cross-validation via partial least square (PLS) regression analysis assigned variable importance in projection (VIP) scores to each taxon to identify key MAG(s) (i.e., critical for these correlations) within each emergent subcommunity (see **Fig. 1A** inset), in effect linking member taxa with host health. We generalized this cross-validated PLS-VIP approach to all emergent sub-communities, using the consensus of multiple statistical techniques, cross-validation methods, and manual inspection (see “*WGCNA analysis*” in **Methods**) to identify key MAGs inferred to drive ecosystem changes, hereafter named “PLS-VIP MAGs”. Only MAGs from male mice networks met the strict criteria of the PLS-VIP MAGs (sex-specific taxon and functional changes are detailed elsewhere^24^).

**Fig. 1.**
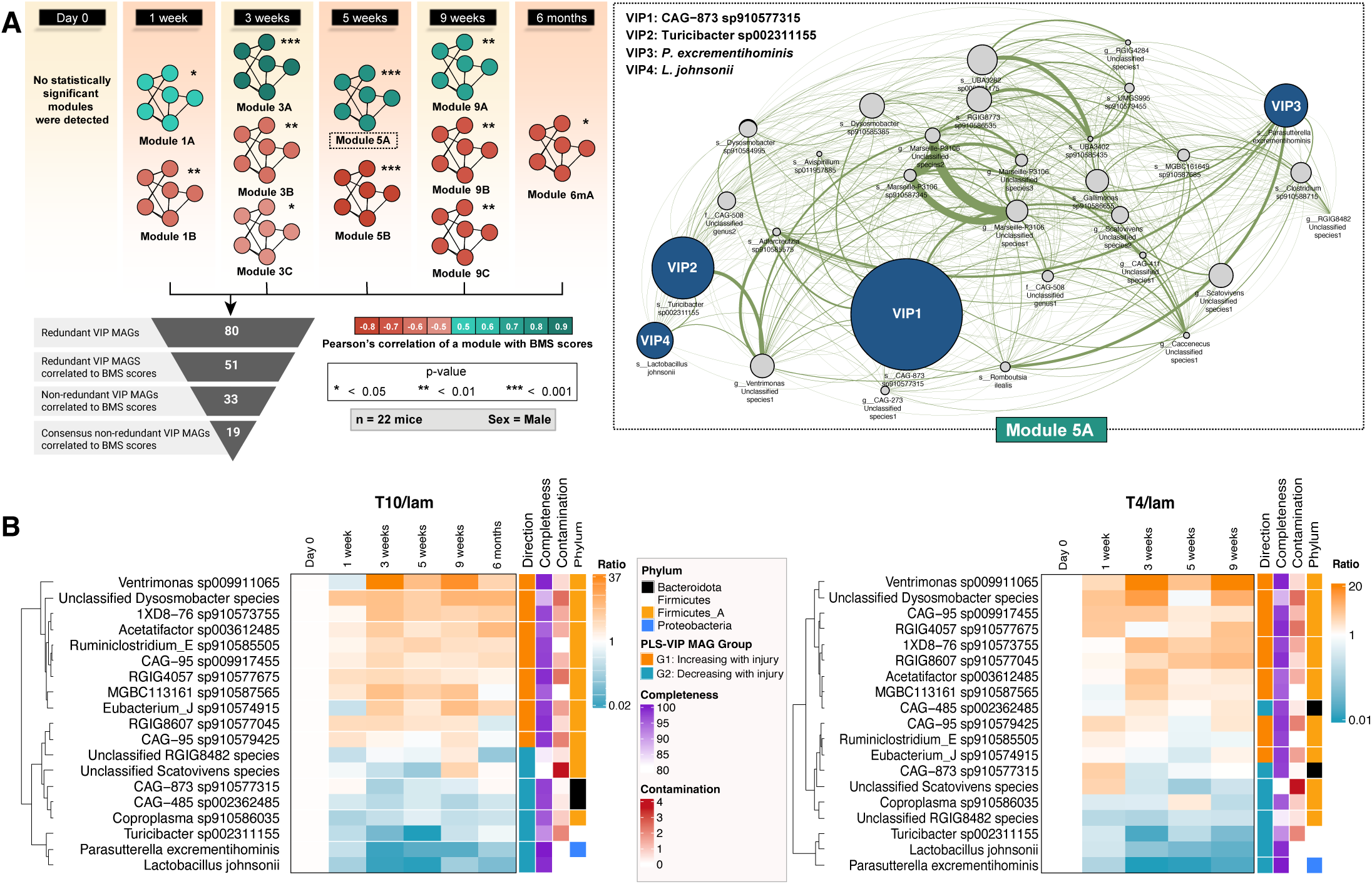
Eco-systems biology distillation of interacting microbiota into host-impacting drivers. **(A)** A schematic of statistically significant sub-communities (in shades of green and red) at each timepoint (see “*WGCNA Analysis*” in **Methods** and **Fig. S1** for details) correlated against 6-months-post-injury BMS score (colored for positive correlations and red for negative correlations and *p*-values as indicated). The modules (subnetworks) are named after the variable time-points: Module 1A = a module emerging at week 1, etc. The inverted pyramid (bottom) shows the number of MAGs filtered at each step of the workflow, distilled to 19 consensus MAGs after accounting for statistical anomalies and normal (i.e., Lam) microbiome drift over time (see “*Inspection of PLS-VIP MAGs*” in **Methods** for details). Inset (right) shows a network visualization of one module (i.e., subnetwork) at 5 weeks after surgery increasing with BMS score. All MAGs within the module are shown in gray except for MAGs highlighted in blue whose VIP scores >=1 and Pearson’s correlation value with BMS score >=0.45. Node sizes are correlated with VIP scores while edge widths are correlated with the strength of correlation between nodes (i.e., MAGs). This module was visualized in Cytoscape v3.7.2. **(B-C)** Heatmaps temporally show the 19 consensus PLS-VIP MAGs that change after NI at T10 and T4 spinal levels, respectively, compared to Lam. The “Ratio” annotation column refers to the ratio of median fold change at an injury level divided by median fold change at Lam to account for normal (i.e., Lam) microbiome drift over the course of the study (see “*Fold-change calculation for visualizing the PLS-VIP MAGs*” in **Methods** for details). G1: Group 1; G2: Group 2.

Application of this workflow to MB6GC MAGs derived from mice with or without NI revealed the following. For non-perturbed gut ecosystems in healthy mice with intact spinal cords, there were no emergent sub-communities at Day 0, confirming similar baseline microbial communities. However, subsequent time points revealed multiple emerging sub-communities after NI (**Fig. 1A**). In total, 19 consensus PLS-VIP MAGs were identified that likely control ecosystem dynamics and function after disrupting spinal cord-gut crosstalk (**Fig. 1**). Of these 19 PLS-VIP MAGs (**Fig. 1B-C**), 11 MAGs from the Firmicutes_A phylum increased in abundance (Group 1 MAGs), whereas 8 MAGs from 4 separate phyla decreased (Group 2 MAGs). Six of the 19 MAGs had strong associations with health or disease states based on prior studies, and of these six MAGs, five decreased in our data (*Lactobacillus Johnsonii*, Turicibacter sp002311155, *Parasutterella excrementihominis*, CAG-485 sp002362485, and CAG-873 sp910577315) and one increased (Acetatifactor sp003612485) after lesioning the spinal cord. Specifically, for these 6 “known” MAGs, published data indicate that these microbes can significantly affect host physiology: (i) *L. johnsonii* enhances gut barrier integrity^39^ and gut mucosal immunity^40^, and reduces plasma total cholesterol levels^41^; (ii) Turicibacter sp002311155 was reduced after murine lung cancer in mice^42^ and strains of the Turicibacter genus differentially modify bile acids and lipid metabolism^43^; (iii) *P. excrementihominis,* a core component of human and murine microbiomes with roles in bile acid maintenance and in cholesterol and tryptophan metabolism, is reduced by high-fat-diet^44^; (iv) CAG-485 sp002362485 (recently proposed as “*Sangeribacter muris*” and biochemically characterized in^45^) is one of the most dominant and prevalent species in the murine gut and may protect from intestinal colitis^45^; (v) similar to *P. excrementihominis*, CAG-873 sp910577315 decreased after high-fat-diet in mice^46^, whereas (vi) Acetatifactor sp003612485 levels were positively correlated with plasma metabolites implicated in cardiovascular diseases^47^. However, while 6 of the PLS-VIP-identified MAGs have previous ties to health and disease in other model systems, our data-driven analyses not only recovered these, but also identified 13 additional MAGs that could have relevance to mammalian host health. Notably, these 19 species-level MAGs were critical to “predict” NI (BMS scores) – due not only to individual abundance changes (perhaps captured by standard pairwise enrichment assessments), but also adding in two more layers of conservative assessment that take into account microbe-microbe co-variation and leave-one-out cross-validation.

### Inferring microbial community metabolic output driven by key network-emergent taxa

Since microbiomes are functionally redundant (i.e., sharing metabolic blueprints) and metabolically connected (i.e, cross-feeding^48^), we assessed how PLS-VIP MAGs might impact host (patho)physiology by analyzing PLS-VIP MAGs’ functional potential in the context of the entire gut microbial community (PLS-VIP MAGs + other community members) metabolism. Instead of relying on changes in single genes, we considered our basic unit to be that of a function, pragmatically represented as genes mapping to metabolic pathways (i.e., expert defined, conserved functional unit of multiple genes). Specifically, we estimated relative abundances of microbial metabolic functions (see **Fig. 2A** and **Methods**) and identified metabolic pathways fulfilling two criteria: (i) those with relative abundances that significantly changed in mice with NI (vs. sham controls) even against community-wide changes (i.e., summing abundances of *all* recovered MAGs encoding these pathways) (**Fig. S2-3**), and (ii) those that were differentially encoded between PLS-VIP MAG groups that increased or decreased in mice with NI as determined by a statistical enrichment analysis (see **Fig. S4** and **Methods**). To identify PLS-VIP MAG-dependent pathway-level changes and minimize pathways functionally compensated by other community members, only pathways meeting both criteria were analyzed further.

**Fig. 2.**
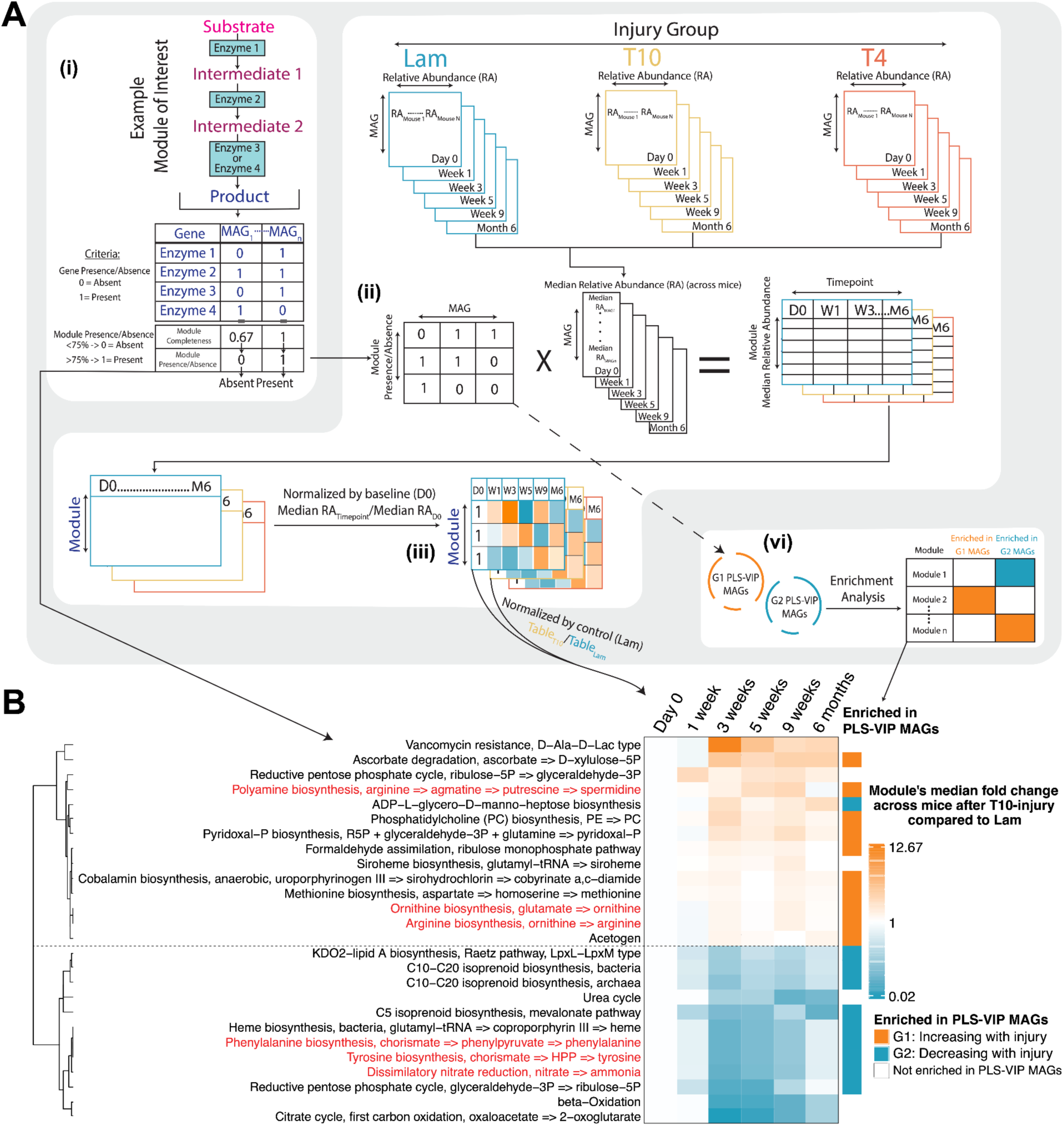
Key metabolic pathways influenced by the spinal cord-gut axis. (**A**) A schematic detailing the workflow developed to determine differentially abundant PLS-VIP MAGs-driven metabolic pathways, including (i) calculating presence/absence of each pathway (i.e., KEGG modules), (ii) integrating the MAG abundance data to calculate the median relative abundance of each pathway, (iii) calculating the module’s fold-change of median relative abundance per injury and across injuries, and (iv) the functional enrichment analysis performed on modules across the two groups of PLS-VIP MAGs. NI at T10 and T4 spinal levels, partially or completely, disrupts neural control over the gut. **(B)** Heatmap representing differentially abundant metabolic pathways at the community level (i.e., all de-replicated MAGs) and their enrichment in the two groups of PLS-VIP MAGs caused by NI. Heatmap values indicate the ratio of median fold changes at T10 divided by median fold change at Lam to account for normal (i.e., Lam) microbiome drift over time in male mice (see “*Fold-change calculation for visualizing pathways*” in **Methods** for more details and **Fig. S2-3** for the complete representation of all pathways following T10 and T4 spinal lesions for male and female mice). Pathways annotated in orange are enriched in PLS-VIP MAGs that increase after NI while those in teal are enriched in PLS-VIP MAGs that decrease. Red text highlights pathways discussed in this report that are thought to be involved in microbial nitrogen metabolism and host nitrogen balance. G1: Group 1; G2: Group 2.

Identifying PLS-VIP MAG-dependent pathways proceeded as follows. *First*, Anvi’o^49^ was used to functionally annotate (see “*Anvi’o pathway identification”* in **Methods**) all 263 MAGs against the KEGG database and organize these annotations (KEGG Ortholog [KO] numbers) into hierarchical KEGG modules that represent partial or complete metabolic pathways. *Second*, module completeness (*sensu*^50^, see **Methods**) was estimated per MAG from the constituent KEGG genes present in it (**Fig. S4A**). *Third*, using abundances of complete modules, a time-resolved, whole-community differential analysis was performed (see “*Fold-change calculation for visualizing pathways”* in **Methods**). This analysis revealed that 26 modules (**Fig. 2B**), as defined by hierarchical clustering (**Fig. S2A**; see all modules in **Fig. S2-3**), encompassed the largest observed module changes in mice with NI, regardless of spinal lesion level or sex. Thus, these 26 modules represent *whole-community* metabolic changes. *Finally,* and independently, a generalized linear model^51^ within Anvi’o (see “*Pathway enrichment analysis with Anvi’o*” in **Methods**) was used to quantify the enrichment of each KEGG module across Group 1 (increasing following NI) and 2 (decreasing) PLS-VIP MAGs. This enrichment analysis identified 52 modules as differentially encoded by the two PLS-VIP MAG groups (**Fig. S4B**).

The intersection of these 52 PLS-VIP MAG-level and 26 whole community-level findings identified 19 complete VIP-driven modules (**Fig. 2B**) that represent the largest abundance-based microbial metabolic changes in mice with NI driven by PLS-VIP MAGs and effectively “ranked” by impact and change direction. We hereafter refer to these modules as “Community VIP (cVIP)-driven modules”. Of these, six modules (**Fig. 2B**) belong to interconnected themes of microbial metabolism critical for host nitrogen metabolism, namely ammonia/urea and amino acid/polyamine metabolisms. Because mammalian physiology related to health/disease is typically correlated with nitrogen balance^36^, where net nitrogen gain and loss are associated with health and disease, respectively, we sought to explore these six modules as most system-relevant.

### Gut microbial nitrogen metabolism disruption by neurological impairment

Among the 19 cVIP-driven modules, one module involved in bacterial fixation of inorganic nitrogen (dissimilatory nitrate reduction, via nitrite, to *ammonia*, hereafter DNRA; KEGG module M00530) was significantly reduced in mice with NI (**Fig. 2B**) and was solely encoded by *P. excrementihominis* among the PLS-VIP MAGs (**Fig. S4**). DNRA enables the retention of scarce nitrogen as soluble ammonia in organic-rich and anoxic ecosystems^52^ and is considered a major pathway for nitrate reduction by bacteria in rumen and human intestine^53–56^.

In mice with NI, we also observed a significant reduction in the bacterial organic nitrogen pathway for urea nitrogen salvaging (UNS) involved in degrading *urea* to ammonia via urease genes. UNS by gut microbes likely has an appreciable role in mammalian nitrogen metabolism^57,58^, albeit typically treated as a “black box”. Indeed, mammals cannot acquire or “fix” nitrogen from the atmosphere like microbes, so they obtain it mostly from dietary proteins, which they then catabolize into nitrogen-rich compounds like amino acids and ammonia. Although the liver is the primary site where protein is metabolized and ammonia detoxified, intestinal bacteria might also participate in nitrogen metabolism as previously suggested and as evidenced from our data as described below.

In our system, we inspected urease-encoding bacteria by manually reconstructing the urease pathway up from single-gene annotations (*sensu*^59^; it is absent from the KEGG hierarchy), and then genome-contextualizing the pathway (as above for cVIP-driven modules). This revealed that, among PLS-VIP MAGs, 100% of the urease pathway was encoded by only CAG-485 sp002362485 and CAG-873 sp910577315. These two MAGs were highly dominant as their median relative abundances represented >35% or ∼16-20% of the microbial community in healthy (i.e., Lam) or male mice with NI, respectively (**Fig. 3A-B**). Further, only four other MAGs, collectively accounting for <1% of the entire microbial community, encoded this complete pathway (**Fig. 3B**). Notably, other MAGs from the same family of the two urease-encoding PLS-VIP MAGs (Muribaculaceae) lacked urease genes (**Fig. 3A**). This suggests relatively recent functional trait evolution in CAG-485 sp002362485 and CAG-873 sp910577315 ecotypes, which would be missed using marker-gene predictions^60^. The relative abundance of these two MAGs was less prominent in healthy female mice (baseline <20% median relative abundance) and less affected by NI. Instead, other non-urease-encoding Muribaculaceae (i.e., Duncaneilla MAGs) dominated the gut of healthy female mice and were significantly decreased in female mice with NI (**Fig. 3B**). These data suggest sex-specific species-driven microbiota-host metabolic interactions, which impacts generalizability of single-sex experiments seeking to assess microbiome impacts.

**Fig. 3.**
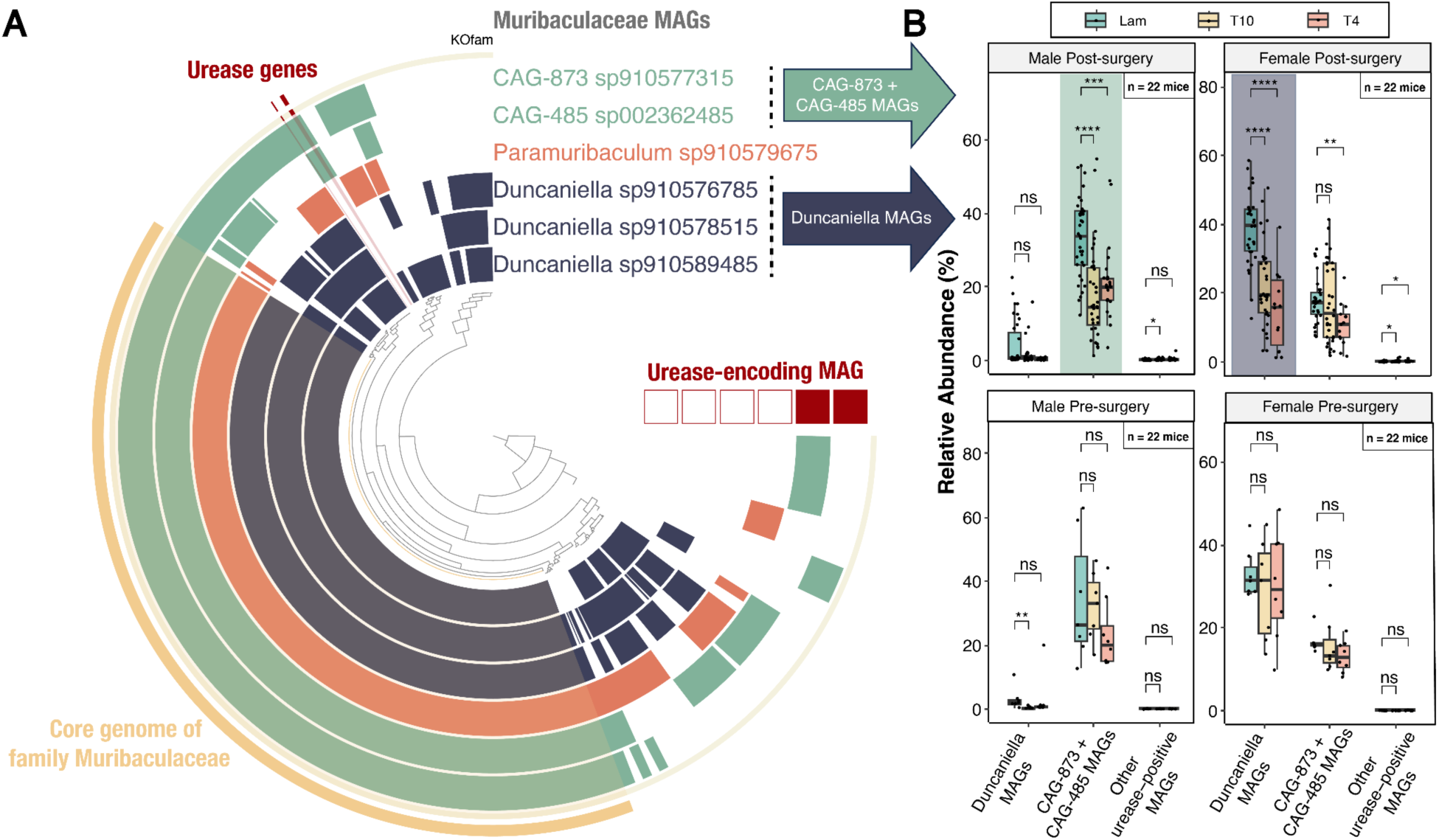
Urease-encoding microbes influenced by the spinal cord-gut axis. (A) Comparative genomics of urease-encoding MAGs within the Muribaculaceae family. MAGs belonging to the same genus are color-coded and MAGs encoding urease genes are indicated by filled red boxes. (**B**) Boxplots (with the 25th and 75th percentile ranges) of aggregate relative abundances of Muribaculaceae MAGs after (top) and before (bottom) NI in male and female mice. Peak changes are highlighted with color-shaded boxes for differentially abundant taxa in male (CAG-873 and CAG-485 MAGs) and female (Duncaniella MAGs) mice. Statistically significant differences in microbial relative abundances between treatments are indicated by the *p*-values from Wilcoxon tests. ****: *p* <= 0.0001, ***: *p* <= 0.001, **: *p* <= 0.01, *: *p* <= 0.05, ns: *p* > 0.05.

Finally, our data revealed significant spinal cord-dependent changes in cVIP-driven modules encoding *amino acids* and *polyamines*. cVIP-driven modules encoding the amino acids tyrosine and phenylalanine, rather than tryptophan^34^, were significantly reduced (**Fig. 2B**, **Fig. 4**). Among PLS-VIP MAGs, only *P. excrementihominis* encoded 100% of these pathways (**Fig. S4**). Conversely, other cVIP-driven modules increased significantly in NI mice. Those modules encode *polyamine* precursors – the amino acids arginine and ornithine – as well as polyamines themselves (**Fig. 2B**, **Fig. 4**). Taxonomically, these biosynthesis pathways were well-represented in Group 1 PLS-VIP MAGs (increasing after NI; 2/11 for polyamine, 11/11 for arginine, and 10/11 for ornithine) but were sparse in Group 2 PLS-VIP MAGs (decreasing after NI; 0/8 for polyamine, and 2/8 for arginine and ornithine) (**Fig. S4**).

**Fig. 4.**
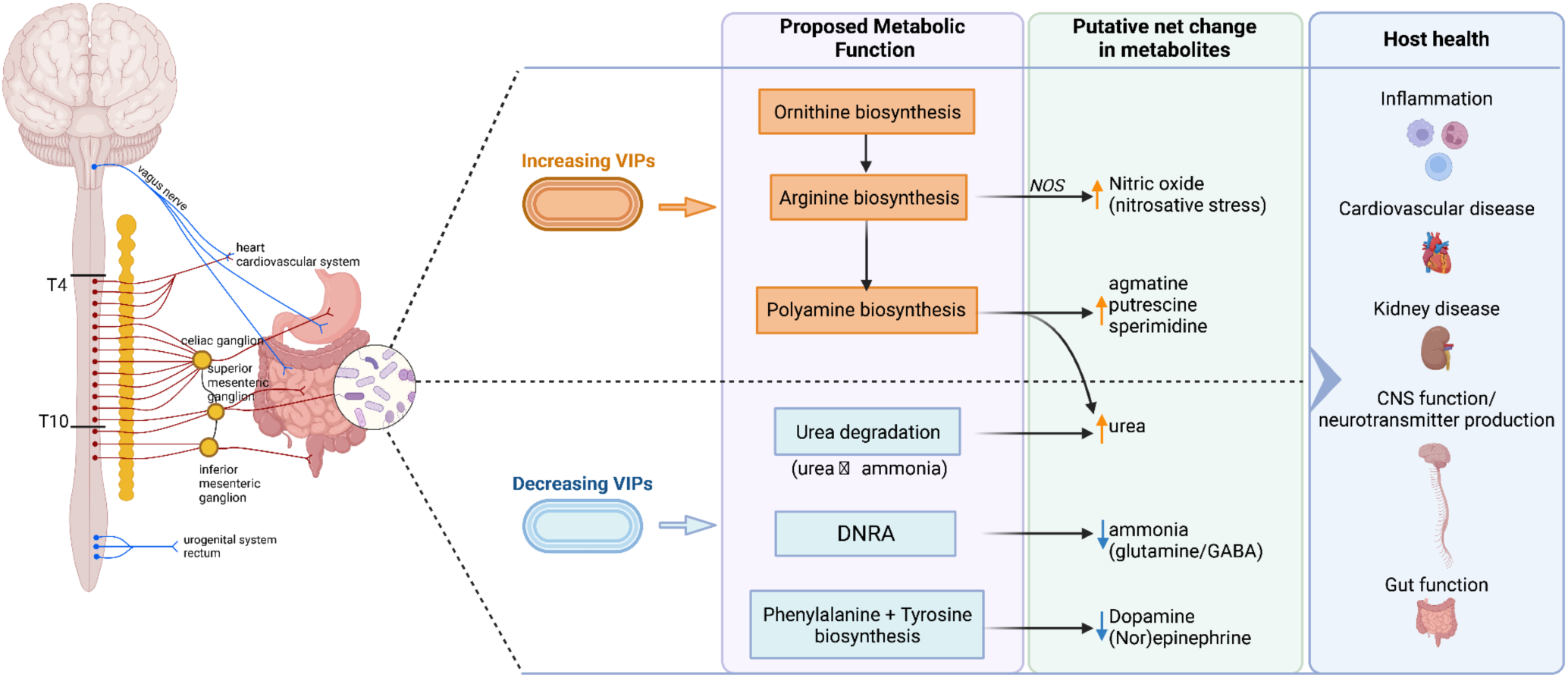
Spinal cord-gut axis impacted microbiota metabolic pathways and their potential effects on host health. Schematic of the spinal cord-gut axis and predicted metabolic functions associated with increasing or decreasing PLS-VIP MAGs following NI. For example (see main text for expanded discussions), PLS-VIP MAGs increasing in abundance (orange) following NI result in an increase in ornithine, arginine, and polyamine biosynthesis. These, in turn, can negatively affect host health via nitrosative stress (i.e., inflammation). Conversely, decreasing PLS-VIP MAGs alter ammonia and urea metabolisms, as well as phenylalanine and tyrosine biosynthesis. A reduction in these MAGs after NI could potentially affect neurotransmitter (e.g., dopaminergic, adrenergic, and GABAergic pathways) availability required for various CNS functions.

## Discussion

This work provides a scalable genome-resolved, functionally contextualized eco-systems biology analytical framework that takes another step away from simple (single-gene/genome/feature) correlations, which fail to consistently infer microbe-host interactions in complex system like the gut^61^, and towards inferred causation and/or mechanism between community-contextualized microbes and host health. The first step in this workflow scales up and semi-automates eco-systems biology approaches developed in another complex ecosystem (i.e., the oceans^37^) to organize taxa or metabolic functions to assess interacting subsets, evaluate their predictive power of ecosystem properties (e.g. carbon flux in the oceans or disease status in mammals), and cross-validate them to assess the specific taxa or functions that are most important to these predictions. Notably, this workflow newly applies this eco-systems biology approach to longitudinal datasets by controlling for temporal dependencies – a critical advance for time-resolved microbiome studies. The second step in our workflow was to establish a mechanism for robustness (i.e., consistency of each taxon’s temporal abundance patterns) by curating identified taxa (i.e., PLS-VIP MAGs) and removing those that were redundant across longitudinally differentiated networks. Finally, the third advance in this workflow was to mechanistically and functionally contextualize changes in consensus PLS-VIP MAGs. This was done by leveraging genome-resolved “complete”-pathway functional signals from multiple data streams that together considered consensus PLS-VIP MAG-based functional enrichment analysis in context of whole-community-based functions. Together, these analyses reduce reliance on pairwise statistical inferences to focus instead on identifying key taxa and complete pathways within community-wide traits and modeled interactions.

Applied to a large-scale dataset from healthy or NI mice, our workflow revealed 19 key whole-community-stabilizing gut bacterial members (i.e., PLS-VIP MAGs; **Fig. 1**) that are influenced by NI and identified 19 microbial metabolic pathways that are modeled to drive microbial ecosystem dynamics in NI mouse guts (**Fig. 2B**). We highlight six of these pathways together with an *ad hoc* reconstructed UNS pathway – all centered around nitrogen metabolism – that suggest and provide specific mechanistic hypotheses for how gut microbiota play a larger role in influencing host nitrogen balance than previously recognized. We discuss here two specific clinically relevant examples related to (i) ammonia biosynthesis and urea degradation pathways, and (ii) altered amino acids and polyamines biosynthesis.

Towards the former nitrogen metabolism impact, NI significantly reduced the abundance of specific dominant gut bacterial MAGs/species previously underappreciated for their role in ammonia biosynthesis and urea degradation pathways (**Fig. 3**), presumably reducing microbially-derived ammonia and elevating urea buildup in the host. The effect of urease-positive bacteria on ammonia levels is supported by decades-long observations^62^ and was recently mechanistically validated and shown to have strong ties to host health^59^, where reduction in bacterium-derived ammonia induced depressive-like behaviors and ammonia-mediated, glutamine-dependent dysfunctional neurotransmission in a (male) mouse depression/stress model^59^. Critically, the brain is the only implicated regulator in this literature. However, based on our data, system-wide changes (e.g., dysfunctional neurotransmission) are expected whenever there is spinal cord dysfunction (likely occurs in virtually all neurological diseases^27–33^) as maintaining organismal homeostasis requires sympathetic nervous system functioning that originates in the spinal cord. Since the spinal cord collects most peripheral tissue sensory information (either directly or later integrated), any nervous system perturbation will impair neurotransmission through the spinal cord and its subsequent control over systemic organs, including the gut. Thus, traumatic spinal cord injury (SCI), through changing microbiota, could alter host health. An example of such alterations is the NI-induced microbiota-mediated ammonia reduction and its subsequent urea accumulation as inferred from our data. Post-SCI uremia (urea buildup) is a common comorbidity in SCI patients that, so far, has been solely attributed to paralysis-dependent muscle catabolism and impaired renal function (based on male rat data^63^). Our data suggest that chronic reduction of gut urease-positive bacteria contribute to that comorbidity, which could also explain why even aggressive nutritional intervention^64,65^ does not allow SCI patients to overcome negative nitrogen balance.

As discussed above, we show that the reduction in abundance of urease-positive bacteria was more pronounced in male mice compared to female mice as the two most abundant species in male mice (accounting for 35% of the baseline community) were urease-positive. Notably, literature reporting ammonia reduction^59^ and urea accumulation^63^ (similar to inferences from our data) used male mice/rats among studies predominately using female mice^5^. We posit that lower baseline levels of urease-positive bacteria in female mice (evidenced by the dominance of urease-negative members from the same family) indicate their lower dependency on the gut microbiome as an ammonia source, compared to males. Consistent with this are data showing that the rate of ammonia generation and recycling in the kidney are greater in female compared to male mice^66,67^. This highlights the importance of controlling for sex in microbiome studies and generates data-driven hypotheses for future experimental validation. Further, we uncover such results that explain cross-disciplinary observations using only a hypothesis-free computational approach, which pinpointed the exact genomes driving systems-level functional changes instead of relying on low-resolution analyses (e.g., only up to family-level predictions of urease-positive bacteria^59^ with little ecological consistency; **Fig. 3**) or on well-characterized taxa that are either rare or transient (i.e., not part of the core microbiome)^59,68^.

Towards the latter nitrogen metabolism impact, the abundance of gut bacterial MAGs encoding aromatic and non-aromatic amino acids and polyamines also was significantly changed in NI mice. First, biosynthesis of aromatic amino acids (tyrosine and phenylalanine) was inferred to decrease following NI. These gut-bacteria-produced amino acids can enter circulation and bypass blood-brain or blood-spinal cord barriers to be precursors for neurotransmitters (dopamine and nor/epinephrine) critical for normal neurological function^69^. Second, biosynthetic pathways for the non-aromatic, semi-essential amino acid arginine and for polyamines increased in NI mice. Because arginine is used as a precursor for synthesizing polyamines and proteins^70^, an increase in gut microbial arginine biosynthesis, or its precursor ornithine, might benefit the host^71,72^. However, literature from murine SCI models shows that an arginine boost may promote inflammation and tissue injury (e.g., neurological recovery was impaired if mice received arginine supplements and neurological function improved if arginine was depleted^73,74^). Similarly, although polyamines are critical for cell growth and viability in mammals^72^, they have been shown to be elevated in serum of uremic patients^75^. Given this conflicting literature and our new findings, we posit that (i) elevated polyamine levels contribute to urea accumulation as the polyamine biosynthetic process releases urea and creates agmatine and/or putrescine (**Fig. 4**), and (ii) the two products (agmatine and putrescine) could play both beneficial and harmful roles in the context of disease or injury to the CNS^76,77^.

Based on the data presented, we propose that increased blood urea associated with NI is due to microbial shifts resulting from disrupted spinal cord-gut communication. Indeed, we see shifts that include (i) a reduction of bacteria encoding urease (particularly in males) and neurotransmitter precursors, and (ii) an increase in bacteria encoding health-critical amino acids and polyamines. Shifting from pre-dysbiotic microbes synthesizing ammonia as their nitrogen source to dysbiotic microbes synthesizing amino acids is consistent with previous reports showing similar metabolic patterns in Clostridia-dominated gut microbiomes^62^. Consistent with this, our data indicate that all 1l PLS-VIP MAGs that increase in NI mice guts belong to the class Clostridia, as opposed to only 3 out of the 8 PLS-VIP MAGs decreasing in these same mice.

Finally, our work shows that understanding community context and interactions is essential when making inferences about local and systemic effects of the gut microbiome (e.g., urea and ammonia balance in the host). The specific example to demonstrate this is the PLS-VIP MAGs identified here presumably serving as the main drivers of this microbial metabolism, with little to no genetic evidence to suggest compensation by rare community members. The lack of community compensation makes these PLS-VIP MAGs an ideal short-list of probiotic targets for future experimental evaluation.

### Conclusions

Medical microbiome research is calling for systems-level genome-resolved approaches that help uncover mechanistic insights to translate the microbiome in health and disease^13,15–17^. This report demonstrates that spinal cord-mediated alterations in gut microbial metabolic potential can significantly impact host nitrogen metabolism. These findings were uncovered using our improved, hypothesis-free, data-driven workflow, which required no *a priori* assumptions or knowledge. By leveraging scalable, genome-informed analytics, we identified a set of microbial taxa, termed PLS-VIP MAGs, that best predict host physiological state and drive the community nitrogen metabolism. While some PLS-VIP MAGs, like *L. johnsonii*, have known associations with neurological outcomes, others are newly predicted to influence host metabolism despite being rarely linked to neurological disorders or considered under the influence of spinal sympathetic networks. If validated in relevant neurological disease models, perhaps a combination of PLS-VIP MAGs could represent broadly impactful probiotics, given their demonstrated metabolic functions (shown here) and ubiquity within the core gut microbiome^78,79^. Such validation experiments represent the next grand challenge as, critically, they must recapitulate community context, such as cross-feeding partners and competing niches, that is inherent in natural microbiomes and captured in our now community-aware analytics. Success in this would empower microbial therapeutics to restore nitrogen balance and alleviate characteristic CNS disease symptoms where spinal cord-gut communication is impaired, including stroke, spinal injury, multiple sclerosis, and myriad neurodegenerative diseases^64,65^. As a step towards this, experiments guided by these approaches now show that restoring one PLS-VIP MAGs can prevent the onset of metabolic and immune dysfunction caused by SCI^24^.

More broadly, however, this report provides a roadmap for addressing a grand challenge in microbiome science whereby microbiota and their functions are more mechanistically linked with their environmental^1,80^ and/or host^14,81,82^ contexts. Given the challenges of *in-vivo*-representative synthetic communities experiments^83^ (e.g., including all interacting partners), we query the natural community interactions across different ‘environmental’ conditions. We do this by shifting from global pairwise predictions about microbial functions based on conserved, slow-evolving marker genes (e.g., 16S rRNA genes) – which are likely evolutionarily out of sync with many ecological traits^5–14^ – towards genome-, population-structure-^12^ and community-informed functional predictions. In this way, our data-driven discovery process can better predict the fast-evolving, niche-defining traits that structure ecosystems and yet often vary significantly within genera and species^10,14^. Such analytical advances, alongside improved community metabolic modeling approaches to assess unintended consequences^84,85^, will expedite the development of systems-level knowledge, which traditionally requires years to achieve but is critical to more fully realize the potential for microbiome engineering as a precise and context-specific therapeutic for personalized medicine.

---

**Fig. S1.**
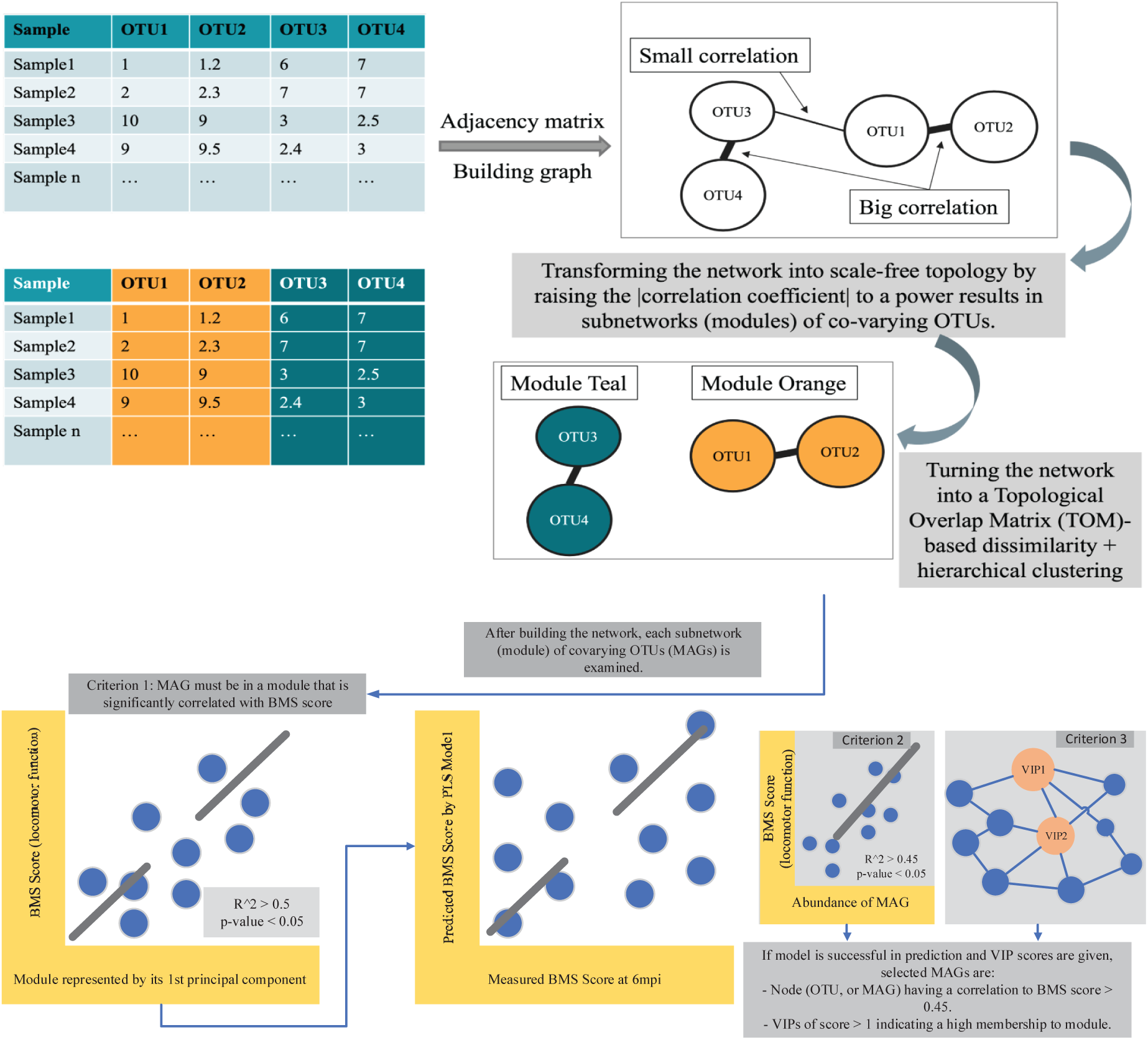
Schematic showing WGCNA analysis to identify key MAGs relevant to neurological outcome after a break in spinal cord-gut communication. MAG abundance-based correlation networks were built separately for male and female samples for each timepoint. The workflow summarizes the steps followed for each of these individual networks (see “*WGCNA Analysis*” in **Methods** for details). First, a graph was built from MAGs (as operational taxonomic units; OTUs) relative abundances and the edges in this network were iteratively transformed (by a power-law function) to meet the scale-free topology assumption for this network. Subnetworks (or modules) from each network were then identified and individually examined across three criteria to identify PLS-VIP MAGs. First, the subnetwork is significantly correlated (correlation coefficient of >=|0.5|, p-value <0.05) to BMS score (as a marker of injury severity). Second, cross-validated partial least squares (PLS) regression between measured and predicted BMS scores for each subnetwork identified the MAG as a key taxon driving the change in this subcommunity by the means of variable importance in projection (VIP) designation (VIP score >= 1). Third, the PLS-VIP MAG is directly or inversely correlated to BMS score using a standard correlation analysis (Pearson’s correlation to BMS score >=|0.45|). Finally, consensus PLS-VIP MAGs were further selected based on manual inspection of statistical anomalies and normal (i.e., Lam) microbiome drift over time (see “*Inspection of PLS-VIP MAGs*” in **Methods** for details).

**Fig. S2.**
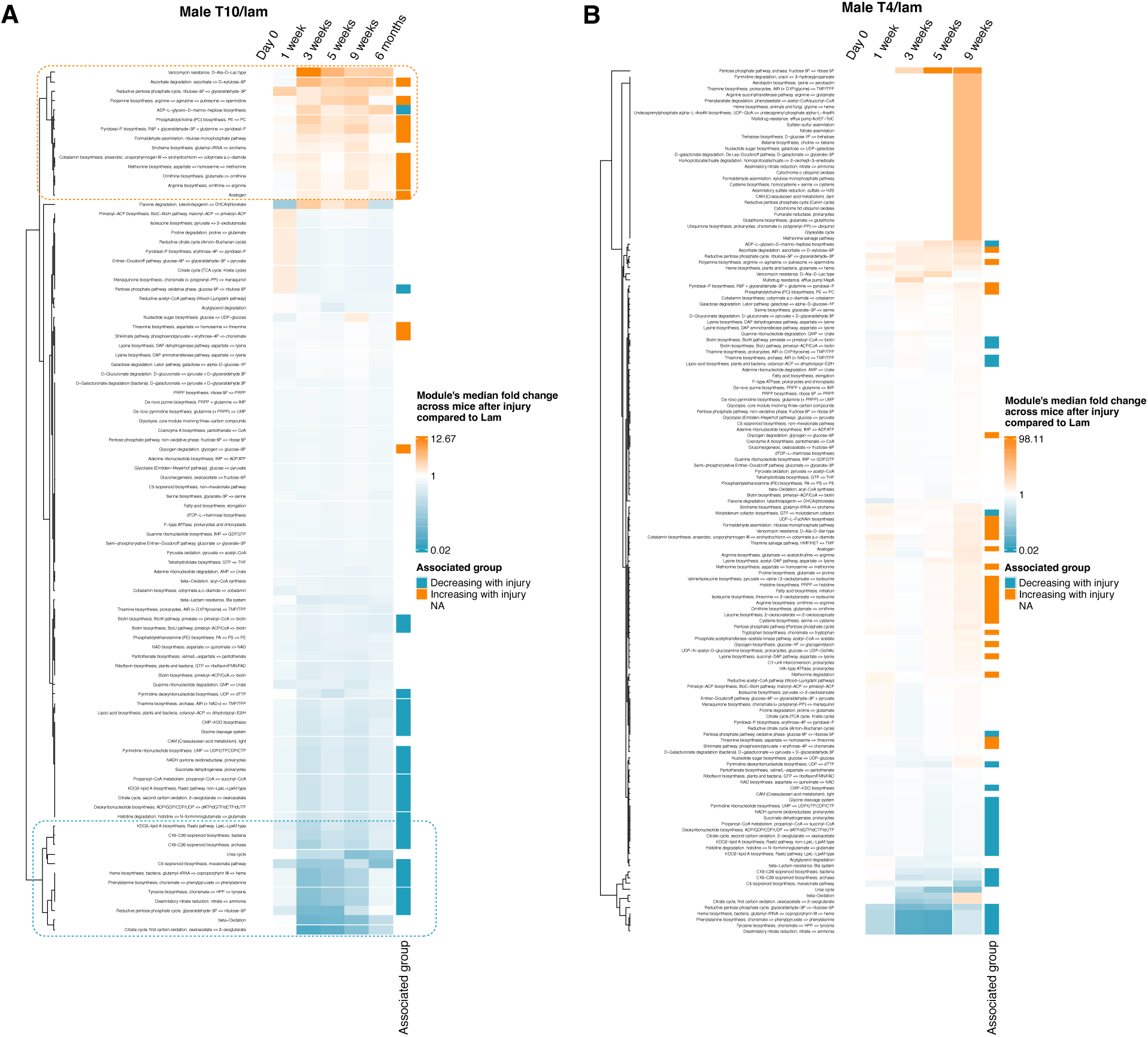
Hierarchical clustering of KEGG pathway changes in male mice. KEGG pathway changes were calculated across the entire community of MAGs to provide an estimate of the overall KEGG module change in the microbiome within injury groups (**A**, T10/lam; **B**, T4/lam) and at different timepoints. Heatmap values indicate the ratio of median fold changes at T10 (**A**) or T4 (**B**) divided by median fold change at Lam to account for normal (i.e., Lam) microbiome drift over time in male mice (see “*Fold-change calculation for visualizing pathways*” in **Methods** for more details). Pathways annotated in orange are enriched in PLS-VIP MAGs that increase after NI while those in teal are enriched in PLS-VIP MAGs that decrease. These modules were compared with the VIP modules from **Fig. S4B** to identify “Community VIP-driven modules” or cVIP-driven modules. The 26 modules within the top and bottom cluster of heatmap (see boxed regions in panel A) are shown in Fig. 2B, which represent the highest changes in T10 (panel A) and encompass all changes in T10-T4 in male and female mice data (see **Fig. S3**).

**Fig. S3.**
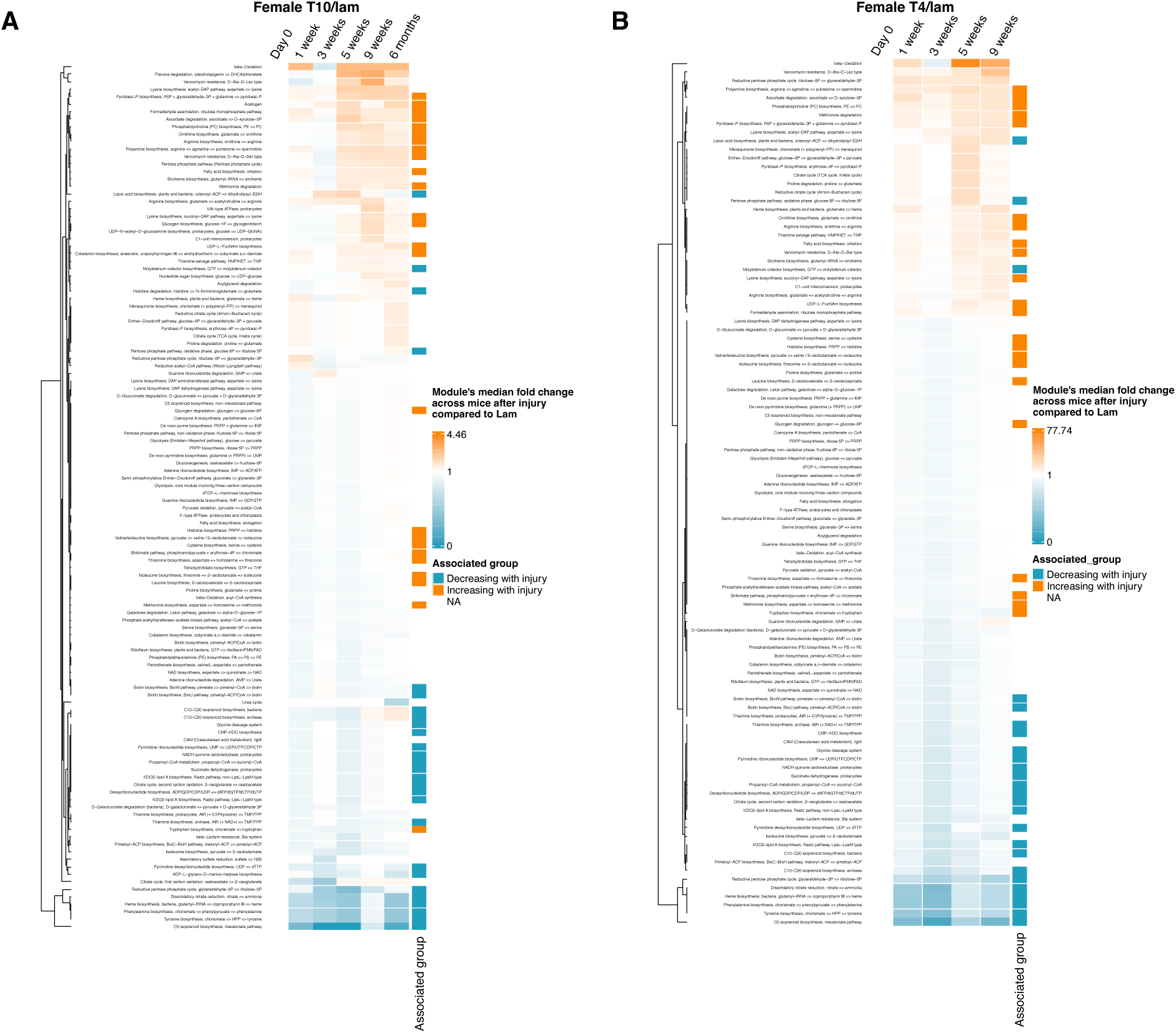
Hierarchical clustering of KEGG pathway changes in female mice. KEGG pathway changes were calculated across the entire community of MAGs to provide an estimate of the overall KEGG module change in the microbiome within injury groups (**A**, T10/lam; **B**, T4/lam) and at different timepoints. Heatmap values indicate the ratio of median fold changes at T10 (**A**) or T4 (**B**) divided by median fold change at Lam to account for normal (i.e., Lam) microbiome drift over time in female mice (see “*Fold-change calculation for visualizing pathways*” in **Methods** for more details). Pathways annotated in orange are enriched in PLS-VIP MAGs that increase after NI while those in teal are enriched in PLS-VIP MAGs that decrease.

**Fig. S4.**
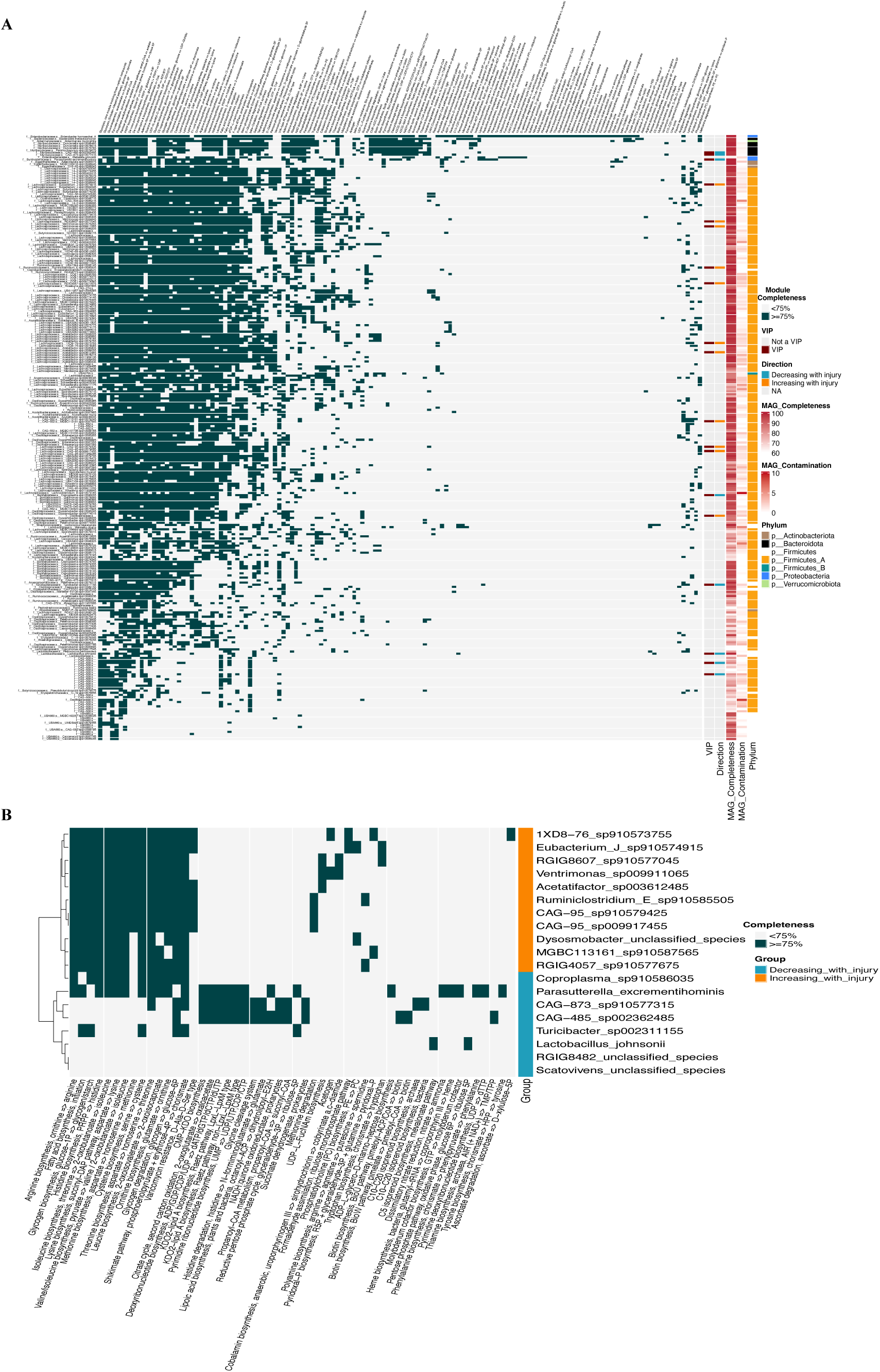
Heatmaps of KEGG modules presence-absence data of all MAGs and of PLS-VIP MAGs that change after a break in gut-spinal cord communication. **(A)** KEGG modules encoded by all 263 de-replicated MAGs. MAGs are annotated with relevant information (e.g., contamination, completeness, phylum, and whether it is a PLS-VIP MAG). **(B)** Significantly enriched KEGG modules in one of the two groups of PLS-VIP MAGs (see “*Pathway enrichment analysis with Anvi’o*” in **Methods**). For (**A-B**), a module is considered encoded by a MAG if the MAG encodes 75% or more of the module’s constituent genes.

## Methods

### Animals and spinal cord injury

All surgical and post-operative care procedures were performed in accordance with the Ohio State University Institutional Animal Care and Use Committee following protocol described in^24^. Briefly, n= 29 female C57BL/6 mice and n=30 male C57BL/6 mice from Jackson Laboratories (Bar Harbor, Maine) were used in this study. To prevent gut microbial cross-contamination due to co-habitation, all mice were single-housed. Mice were anesthetized with an intraperitoneal cocktail of ketamine (80 mg/kg)/xylazine (10 mg/kg) after which a partial laminectomy was performed at the fourth thoracic spine (T4) or the tenth thoracic spine (T10). In mice modeling SCI, modified #5 Dumont forceps were inserted laterally and held closed for 3 seconds to produce a complete crush spinal cord injury. In total, 20 mice received a complete crush at T4, 20 mice received a complete crush at T10, and 19 mice received laminectomy (Lam) only.

Fecal samples for metagenomic sequencing were aseptically collected pre-injury, 1-week, 2-weeks, 3-weeks, 5-weeks, 9-weeks, and 6-months post-injury, immediately frozen and stored at −80℃.

#### WGCNA analysis (i.e., to identify MAGs Relevant to spinal cord-gut cross-talk)

To identify key MAGs, or species, driving dysbiosis in the microbial community from a healthy state (i.e., Lam mice) to a diseased state (i.e., T10- or T4-injured mice), co-occurrence networks were built with hellinger-transformed MAG abundances using the “decostand” function from the R package “vegan” (https://github.com/vegandevs/vegan). These networks were built under the framework of WGCNA^86^ (R package: https://cran.r-project.org/web/packages/WGCNA/index.html), a systems biology approach used to detect significant associations between genes or taxa and a system variable (BMS-score of mice at 6-months post injury in this study). This continuous variable accounts for incomplete injuries or partial crushing of the spinal cord rather than using a categorical variable such as injury levels. Here, WGCNA was used on male and female cohorts separately, and one network was built per each timepoint. All codes used were deposited in the accompanying Zenodo repository.

#### For building networks

Hellinger-transformed abundance tables were used to build the networks using the biweight midcorrelation method (corType=“bicor”). The values used as soft thresholds to ensure the networks’ free-scale topology were 10, 12, 12, 14, 16, and 18 for pre-injury (i.e., baseline, or Day 0), 1 week, 3 weeks, 5 weeks, 9 weeks, and 6 months post-injury, respectively. The corresponding values for female mice data were 10, 12, 14, 14, 10, 14. Networks were built by the blockwiseModules function in the WGCNA package, requiring the “signed” option for both networkType and TOMType, and a minimum module size of 15.

#### Finding modules significantly correlated to BMS score

For each subnetwork, or module, of covarying MAGs extracted from each network (e.g. per timepoint in males), pairwise Pearson’s correlation coefficients and their corresponding *p*-values were calculated between the module’s first principal component and the BMS score. A module was considered significant if it has a correlation coefficient of |0.5| or larger and a *p*-value of less than 0.05. The modules with the highest correlation scores were examined further. These included two modules at 1 week, three at 3 weeks, two at 5 weeks, three at 9 weeks, and one at 6 months. No modules with significant correlation were found at Day 0, corroborating that there were no baseline pre-injury differences. All significant modules and their correlation coefficients and *p*-values are shown in **Fig. 1A**.

#### PLS-VIP MAGs selection

To identify taxa driving changes per module, we applied partial least squares (PLS) regression to determine the best predictors of BMS score. This was validated by a leave-one-out cross-validation method. All predictors per module were ranked according to their contribution in the PLS regression, which is known as their value importance in projection (VIP). MAGs with larger than |0.45| Pearson’s correlation to BMS score that also had a VIP score larger than 1 were considered important for the PLS prediction and were thus further investigated. We refer to these MAGs as “PLS-VIP MAGs” throughout the manuscript (or “VIPs” for short).

#### Inspection of PLS-VIP MAGs

To identify “consensus” PLS-VIP MAGs, each MAG was tested a) temporally within each treatment (i.e., injury levels) and b) across treatments per each timepoint, both using pairwise wilcoxon tests as described above. Unless otherwise stated (see full rationale for the selection criteria for each MAG in the accompanying Zenodo repository), PLS-VIP MAGs were kept in the “consensus” list if: a) their abundances significantly changed over time in T10 and/or T4 mice compared to baseline (Day0) when testing was constrained by treatment; and b) their abundances significantly changed between Lam (control) and T10 or Lam and T4 at one or more timepoints when testing was constrained by timepoint. In both testing strategies, trends in control samples (e.g., significant changes in Lam over time) and at baseline (e.g., significant changes in T10 or T4 compared to Lam at Day0) were used to inform exclusion from the consensus list.

#### Fold-change calculation for visualizing the PLS-VIP MAGs (Fig. 1B-C)

In each mouse, the MAG’s relative abundance at each timepoint was divided by its relative abundance at Day 0 after introducing small pseudo-counts (both quantities were multiplied by 1000 and added to 1) to avoid division by zero. The median fold-changes across mice was calculated for each taxon per injury and timepoint. The median fold-changes in T10 or T4 were then divided over those in Lam for each taxon and then plotted. The heatmap was constructed with the ComplexHeatmap package^87^ (https://github.com/jokergoo/ComplexHeatmap) and all values were log2-transformed to improve color ranges.

### Anvi’o pathway identification

Using Anvi’o v7.1 (https://github.com/merenlab/anvio) and all de-replicated MAGs as external genomes, the following commands were used with their default parameters on each MAG: anvi-gen-contigs-database, anvi-run-hmms, anvi-run-kegg-kofams, and anvi-estimate-metabolism (with --kegg-output-modes modules, kofam_hits and --matrix-format). This gives identified KEGG modules and KO hits per MAG as well as module completeness and presence-absence matrices per MAG. The default value (75%) for module completeness was used.

### Pathway enrichment analysis with Anvi’o

Using Anvi’o v7.1 and the PLS-VIP MAGs as external genomes, the anvi-compute-metabolic-enrichment function was used to compute enrichment in the 2 groups of PLS-VIP MAGs (i.e., the ones enriched in T10 or T4 compared to Lam and the ones enriched in Lam compared to these two injuries). KEGG modules with an adjusted q-value of less than 0.05 or ones that are present in one group while absent in the other were considered and plotted in **Fig. S4B**.

### Fold-change calculation for visualizing pathways

(Fig. 2B and Fig. S2-3) The abundance of a KEGG module (i.e., pathway) was calculated from the total of the abundances of the MAGs encoding it. First, the median abundance of each MAG across all mice per sex, injury, and timepoint was calculated. Next, this median abundance was multiplied by the presence-absence value of the KEGG Module (1 or zero) determined by Anvi’o as explained above, where a module is considered present if it is at least 75% complete. These quantities (median MAG abundance across mice belonging to the same injury and timepoint, multiplied by the module’s presence/absence value) were then summed for all MAGs to give an estimate of the overall KEGG module’s abundance per sex, injury and timepoint. For each sex, these values at each timepoint, per injury, were then divided by the values at Day0 to calculate the median fold change. Finally, T10 and T4 values were divided by the corresponding Lam values. Heatmaps were constructed with the ComplexHeatmap package and all values were log10-transformed to improve color ranges.

### Data availability

All data produced in this study are publicly available. Curated data and all used codes and databases can be found in the following Zenodo DOI: 10.5281/zenodo.15613780 (Unpublished - preview [here](https://zenodo.org/records/15613780?preview=1&token=eyJhbGciOiJIUzUxMiJ9.eyJpZCI6IjVhMDY0YzgxLTA2ZTgtNGU3Yi1iNjUxLWZjZDBkOTE1ODk5NyIsImRhdGEiOnt9LCJyYW5kb20iOiI1NTAxMDY5ZTBhYWZiOTVkMTY3YTIzMzM5MzQ1YWVlNiJ9.TNCyCSUT2Lpl6qLQcjjdCD3nhGPsIx9VwSIrzLZLari80GJL0Sy39zUQ69dn_6lEKg4gezhchZ8_2h56qhP6ZA)).

## Acknowledgements

We appreciate the help and support from all members of the Popovich and Sullivan laboratories, especially Natalie Solonenko, who participated in data sequencing, and Ami Fofana for her immense help with visual representations. We’d also like to acknowledge the Center of Microbiome Science, Ohio Supercomputer Center, and the Riffomonas educational resource^88^. This work is funded by a National Institutes of Neurological Disorders and Stroke R35 award (grant no. 1R35NS111582) to P.G.P., The Belford Center for Spinal Cord Injury (P.G.P.), The Ray W. Poppleton Research designated endowment (P.G.P.), a Craig H. Nielsen Foundation Senior Research award (grant no. 890085) to K.A.K, and NSF awards ABI#2149505 and DBI#2022070 to M.B.S.

## Author Contributions

P.G.P., M.B.S., K.A.K., A.A.Z. conceived the project and experimental design. P.G.P., M.B.S., K.A.K., A.A.Z., and M.M. developed the data analysis plan. P.G.P., M.B.S., K.A.K., A.A.Z., M.M., G.J.S., J.D., and J.M.S. helped with data interpretation and writing of the manuscript. M.M. performed the statistical, functional, bioinformatic, and eco-systems biology analyses, and managed and coordinated responsibilities for the manuscript preparation. A.A.Z. guided and supervised statistical, functional, and bioinformatic analyses, performed eco-systems biology analyses, co-developed the WGCNA pipeline, and drafted the manuscript with M.M.. M.M., K.A.K., A.A.Z., and G.J.S. contributed to figure generation and, with P.G.P., contributed to improving illustrations. G.J.S. guided functional analyses. All authors read and reviewed the manuscript and approved it in its final form.

## Declaration of generative AI and AI-assisted technologies

During preparation of this work, Microsoft CoPilot and ChatGPT were used periodically to improve writing or shorten passages. After using it, the authors reviewed and edited the content as needed and take full responsibility for the content of the publication.

